# Dual stimulation of CD40 and 41BB pathways during ex-vivo TIL expansion enhances CD8+ T cell expansion

**DOI:** 10.64898/2026.07.22.740128

**Authors:** Renata Ariza Marques Rossetti, Matthew S. Beatty, Junior Cianne, Johannes R. Ali, Kylie Harris, Ahmed Ramadan, Michael Grant, Elena Martinez Planes, Jyllica Aurelio, Lilit Karapetyan, Ben Creelan, Shari Pilon-Thomas, Patrick Hwu, Vincent C. Luca, Daniel Abate-Daga

## Abstract

**Background:** Tumor-infiltrating lymphocyte (TIL) therapy has demonstrated clinical efficacy in malignant melanoma; however, inefficient ex vivo expansion remains a major limitation. We previously showed that stimulation of tumor-infiltrating B cells via CD40-CD40L axis improves TIL expansion, and that direct activation of the 41BB–41BBL pathway on T cells enhances CD8⁺ T cell outgrowth. We hypothesized that adding simultaneous targeting of both pathways would augment the growth and activity of CD8+ cytotoxic T cells. We conducted a study with the objective of determining the feasibility of dual stimulation with human tumors as justification for a Phase I trial.

**Methods:** CD40L variants were generated by yeast display selection and evaluated for B cell binding and activation. The effects of CD40L variants on TIL expansion were evaluated using tumors derived from standard of care resections using fragment method. Based on these findings, a bi-specific molecule was designed and generated fusing a CD40L variant and 41BB to the N- and C-termini of a trimeric leucine zipper. The effects of the bi-specific molecule (termed CD40L^EPC6^-41BBL) on TIL expansion were evaluated in TIL cultures derived from lung tumor and melanoma fragments. TIL phenotypes were assessed by flow cytometry, including high-dimensional FlowSOM analysis, and tumor reactivity by autologous tumor co-culture assays.

**Results:** Each of our engineered CD40L variants bound B cells and induced CD80/CD86 expression at levels comparable to wild-type CD40L. Supplementation of TIL cultures with CD40L variants increased the success rate of TIL expansion compared to control. We then developed a bi-specific CD40L^EPC6^–41BBL molecule capable of binding to both B and T cells. Addition of CD40L^EPC6^–41BBL significantly increased total TIL yield and improved expansion success rates in both lung tumor and melanoma cultures. In particular, CD40L^EPC6^–41BBL promoted preferential expansion of CD8⁺ T cells. High-dimensional analysis revealed enrichment of CD8⁺ T cell clusters expressing CD39, CD69, TIM3, and CD56 in cultures supplemented with CD40L^EPC6^–41BBL. Furthermore, treated cultures displayed increased frequencies of CD27⁺ CD4⁺ T cells. Functional assessment suggested a trend toward enhanced tumor reactivity in melanoma-derived TIL products expanded with CD40L^EPC6^–41BBL.

**Conclusions:** Simultaneous stimulation of CD40 and 41BB pathways using a novel bi-specific molecule resulted in qualitative and quantitative enhancement of TIL products. These findings support dual targeting of tumor-infiltrating B cells and T cells as a promising strategy to optimize TIL manufacturing for adoptive cell therapy in Phase I trials.

## BACKGROUND

Tumor infiltrating lymphocyte (TIL) therapy today represents the most clinically mature adoptive cellular therapy modality for solid tumors, particularly melanoma.^1^ The therapeutic efficacy of TIL therapy relies on the expansion of tumor-reactive T cells from resected tumor specimens, followed by reinfusion of these cells into the patient.^2^ Despite encouraging clinical outcomes, the generation of TIL products remains challenging, as many tumor samples fail to yield sufficient TIL numbers during ex vivo culture.^3^ Furthermore, expanded TIL products have a high heterogeneity in cellular composition and functional characteristics, which influence clinical efficacy.^4^ Therefore, strategies to improve the expansion of efficient tumor-reactive T cells remain an important area of investigation. TIL manufacture typically involves an initial culture period of tumor fragments in presence of high-dose interleukin-2 (IL-2). Increasing evidence suggests that cells within the tumor microenvironment can influence the quantity and quality of the expanded TIL product.^5^ For example, we and others have demonstrated that tumor-infiltrating B cells can be leveraged to enhance ex vivo TIL expansion through indirect stimulation of T lymphocytes.^6^ ^7^ The ultimate mechanism of this indirect stimulation is not fully understood, but is likely to involve multiple co-stimulatory ligands including CD83 and CD58.^6^ ^8^ Direct co-stimulation of T cells using anti-41BB agonistic antibodies has also proven effective in enhancing the properties of TIL products.^9^ ^10^ Based on these observations, we hypothesized that simultaneous targeting of B cells and T cells through complementary co-stimulatory pathways may further improve TIL expansion. In the present study, we first engineered and characterized novel CD40L variants with improved biochemical properties while maintaining their ability to bind and activate B cells. We then developed a bi-specific protein combining the cognate ligands of CD40 and 41BB (CD40L and 41BBL, respectively), enabling concurrent stimulation of CD40-expressing B cells and 41BB-expressing T cells during TIL culture. Using lung tumor and melanoma samples, we evaluated the effects of this bi-specific protein on TIL expansion, phenotype, and tumor reactivity. Our findings demonstrate that simultaneous targeting of CD40 and 41BB pathways represents a promising strategy to optimize ex vivo TIL manufacturing for adoptive cell therapy.

## METHODS

### Yeast display selection of CD40L variants

The ectodomain of human CD40L was cloned into a modified version of the yeast display vector pCT302 containing a C-terminal HA epitope (YPYDVPDYA), c-Myc epitope (EQKLISEEDL), and Aga2. To generate a mutant library of CD40L, error-prone PCR (GeneMorph II Kit, Agilent Technologies) was used to introduce mutations in the CD40L gene. EBY100 yeast were then electroporated (BTX electroporator) with 10 μg vector and 50 μg of the mutated insert to allow for library assembly via homologous recombination as previously described.^11^ Yeast transformants were recovered in synthetic dextrose media with casamino acids (SDCAA) and the library was displayed through induction with synthetic galactose media with casamino acids (SG-CAA). Several rounds of selections were performed to isolate CD40L variants with enhanced expression/stability. During each round, a positive selection was performed by incubating induced yeast with biotinylated CD40, washing 3x with PBS containing 0.5% BSA, and staining with streptavidin labeled with Alexa Fluor 647.^12^ Binders were then selected using magnetic-activated cell sorting (MACS, Miltenyi Biotec) or flow cytometry. Prior to each round of positive selection with CD40, a negative selection was performed using SA-AF647 alone to eliminate non-specific binders. After selections were completed, sequencing of the enriched yeast revealed an L259F mutation present in all CD40L clones. One clone containing only the L259F mutation, EPC6, was used for the subsequent generation of the CD40L-41BBL fusion protein described below).

### Expression and purification of CD40L^EPC6^ and the CD40L^EPC6^-41BBL fusion proteins

The ectodomains of CD40L or CD40L variants were cloned into the Baculovirus transfer vector pAcGP67a and contained C-terminal 8xHis tags. The ectodomain of CD40 was cloned into pAcGP67a and contained a C-terminal biotinylation acceptor peptide (BAP) and a 6xHis tag. The CD40L^EPC6^-41BBL fusion was also cloned into pAcGP67a and contained an N-terminal CD40L^EPC6^ ectodomain followed by a trimeric GCN4pII leucine zipper plus linker module (previously described for CD40L-tri)^13^, followed by the ectodomain of 41BBL and a 8xHis tag. Each vector was transfected into SF9 insect cells to generate recombinant Baculovirus, and then the amplified virus was used to infect 1L cultures of *T. ni* cells to express the proteins. The infected cultures were then harvested, and proteins were purified from the supernatants using nickel and size exclusion chromatography.

### Thermal denaturation assays

The ectodomains of CD40L and other CD40L variants were diluted to a final concentration of 5 μM in 20 μl HBS buffer (20 mM HEPES pH 7.4, 150 mM NaCl) and then combined with 10 μM (5×) SYPRO Orange (Thermo Fisher Scientific). The mixtures were then equilibrated at room temperature for 30 minutes and samples were analyzed in a 96-well microtiter plate using a StepOnePlus RT–PCR system with software v.2.3 (Applied Biosystems) applying a continuous heating gradient from 25 °C to 99 °C with a step of 1% of temperature per minute.

### Binding-affinity assay

For characterization of CD40L variants’ binding to B cells, PBMC from healthy donors were incubated with the constructs for 20 minutes at 4 °C to allow binding. Cells were then washed and stained with CD19-BV605, CD3-BUV395, HLA-DR-FITC, and His-Tag-Alexa Fluor 647 antibodies. For analysis of CD40L^EPC6^-41BBL binding to T lymphocytes, CD3+ cells were isolated from healthy donor PBMC using the Pan T Cell Isolation Kit (Miltenyi Biotec) and stimulated with Dynabeads magnetic beads (Thermo Fisher Scientific) for 24 hours. T cells were then incubated with CD40L^EPC6^-41BBL for 20 minutes at 4 °C. After washing, T cells were stained with CD3-BUV395, CD4-BUV496, CD8-PECy7, and His-Tag-AlexaFluor647 antibodies. To confirm 41BB expression upon stimulation, a second panel included CD3-BUV395, CD4-BUV496, CD8-PECy7, and 41BB-BV421 antibodies. For all panels, viability was assessed using LIVE/DEAD Near-IR staining. Cells were fixed and acquired on an LSR-II flow cytometer (BD Biosciences), and data were analyzed using FlowJo software (TreeStar). Antibody details are listed in Supplemental Table 1.

### B cell stimulation

For characterization of B cell stimulation by the CD40L, B cells were isolated from healthy donor PBMC using the Pan B Cell Isolation Kit (Miltenyi Biotec) and cultured in presence of CD40L constructs plus an anti-His-Tag antibody (Cell Signaling; 0.5 μg /mL), serving as a crosslinking antibody, for 48 hours at 37 °C. Following stimulation, B cells were stained with CD19-BV605, CD3-BUV395, HLA-DR-FITC, CD80-BV650, CD86-PECy7 and CD40-BV785 antibodies. Viability was assessed using LIVE/DEAD Near-IR staining. Cells were fixed and acquired on an LSR-II flow cytometer (BD Biosciences), and data were analyzed using FlowJo software (TreeStar). Antibody details are listed in Supplemental Table 1.

### TIL culture

Metastatic melanoma, and non-small cell lung cancer (NSCLC) samples were obtained from consenting patients treated at Moffitt Cancer Center. The study was approved by the Advarra IRB (approval numbers MCC20559, and MCC21971). For the initial TIL expansion phase (pre-REP), TILs were expanded for 3-4 weeks from 1–2 mm² melanoma and lung tumor fragments in 24-well plates under the conditions described in the Results section. Culture with 6000 IU/mL IL-2 (Proleukin, Novartis, Emeryville, CA), served as a control condition. TIL culture media consisted of RPMI 1640 supplemented with L-glutamine, 10% heat-inactivated human AB serum, 10 mM HEPES, 100 IU/mL penicillin, 100 ug/mL streptomycin, 50 ug/mL gentamicin, 55 uM 2-mercaptoethanol, and 62.5 mg/mL Amphotericin-B. Half of the media was replaced every 3-4 days or as needed, and wells were sub-cultured when 80% confluent.

### TIL phenotyping

TILs generated during the pre-REP phase from lung tumor and melanoma fragments were stained with CD3-BUV395, CD4-BV785, CD8-PECy7, CD39-FITC, CD69-APC, CD62L-BV421, CD27-BV605, TIM3-BV650, PD-1-Alexa Fluor 700, and CD56-PE antibodies. Viability was assessed using LIVE/DEAD Near-IR dye. Antibody details are provided in Supplemental Table 1. Following surface staining, cells were fixed and acquired on an LSR-II flow cytometer (BD Biosciences), and data were analyzed using FlowJo software (TreeStar). For the high-dimensional flow cytometry analysis, events from each fragment were down-sampled to 5,000 cells for lung tumor samples and 10,000 cells for melanoma samples, then concatenated according to culture condition. These concatenated files were used for dimensionality-reduction analysis (tSNE) and clustering using self-organizing maps (FlowSOM). For each tumor type, a reference map was generated from a concatenated file containing all fragments of that tumor type, irrespective of culture condition, and this map was subsequently applied to the analyses.

### Tumor reactivity assay

Tumor tissues were minced with scalpels and digested mechanically and enzymatically in RPMI 1640 with L-glutamine supplemented with 30 U/mL DNase type IV, 100 µg/mL hyaluronidase type V, 1 mg/mL collagenase type IV, 50 µg/mL gentamicin, 100 IU/mL penicillin, 100 µg/mL streptomycin, and 62.5 mg/mL amphotericin B. Single-cell suspensions were counted, and cryopreserved for later use. For tumor reactivity assays, autologous tumor cells were co-cultured with TILs at a 1:1 ratio in 96-well round-bottom plates for 24 hours. As a control for HLA-dependent reactivity, tumor cells were pre-incubated with anti-human HLA-A, B, C blocking antibody (W6/32, Biolegend; 10 mg/mL) for 1 hour at 37 °C prior co-culture. As a positive control, plates were pre-coated with anti-human CD3 antibody (Clone OKT3, Biolegend; 5 mg/mL). After 24 hours, culture supernatants were collected, and tumor reactivity was quantified by measuring IFN-γ, TNF-α, and Granzyme B secretion using the ELLA automated immunoassay system (ProteinSimple, San Jose, CA).

### Statistical analysis

All statistical analyses were performed using GraphPad Prism software unless otherwise specified. **CD40L mutant analysis:** Binding to B cells was evaluated using ordinary one-way ANOVA. Effects of CD40L stimulation on the percentage of CD80⁺CD86⁺ B cells were assessed using two-way ANOVA with Tukey’s multiple comparisons test and repeated-measures ANOVA with Tukey’s correction. The effect of CD40L addition on TIL expansion success was analyzed using a two-sided Chi-square test. **Fusion molecule analysis:** Binding to B and T cells and the effects of CD40L^EPC6^–41BBL stimulation on CD80⁺CD86⁺ expression were evaluated using unpaired *t*-tests. The impact of CD40L^EPC6^–41BBL on TIL expansion and CD4/CD8 composition was assessed using the Wilcoxon paired test and two-sided Chi-square test. Comparisons of the frequency of CD27⁺ cells within the CD4⁺ T-cell subset were performed using the Mann–Whitney test. A *p* value ≤ 0.05 was considered statistically significant.

## RESULTS

### Selection of stabilized CD40L variants

We previously showed that melanoma-infiltrating B cells express CD40 and respond to stimulation via CD40L^6^. Trimerization of membrane-bound CD40L is important for signaling, whereas soluble versions of CD40L do not robustly signal in the absence of a multimerization scaffold.^13^ To engineer biochemically enhanced CD40L variants for TIL expansion, we used yeast surface display to select for mutations that increased expression, stability, and/or binding to CD40 (**Fig. 1A**). A library of >10^7^ CD40 variants was expressed on the surface of yeast, and several rounds of selection were performed to enrich for clones with improved binding to the recombinant CD40 ectodomain, or to an epitope-specific antibody (c-Myc) that detects surface expression (**Fig 1B**). Sequencing of individual clones revealed a recurrent L259F mutation (**Fig. 1C**), including one variant that included only this mutation (henceforth referred to as EPC6). Structural analysis revealed that the L259F substitution positions a bulky aromatic group in the trimer interface of CD40L (**Fig. 1D**)^14^, and differential scanning fluorimetry (DSF) revealed that the melting temperature of CD40L EPC6 (76.6 °C) was increased by ∼13 °C compared to that of CD40L WT (62.9 °C) (**Fig. 1E**). These data suggest that L259F enforces CD40L trimerization on the surface of yeast cells, which in turn promotes receptor binding.

**Figure 1.**
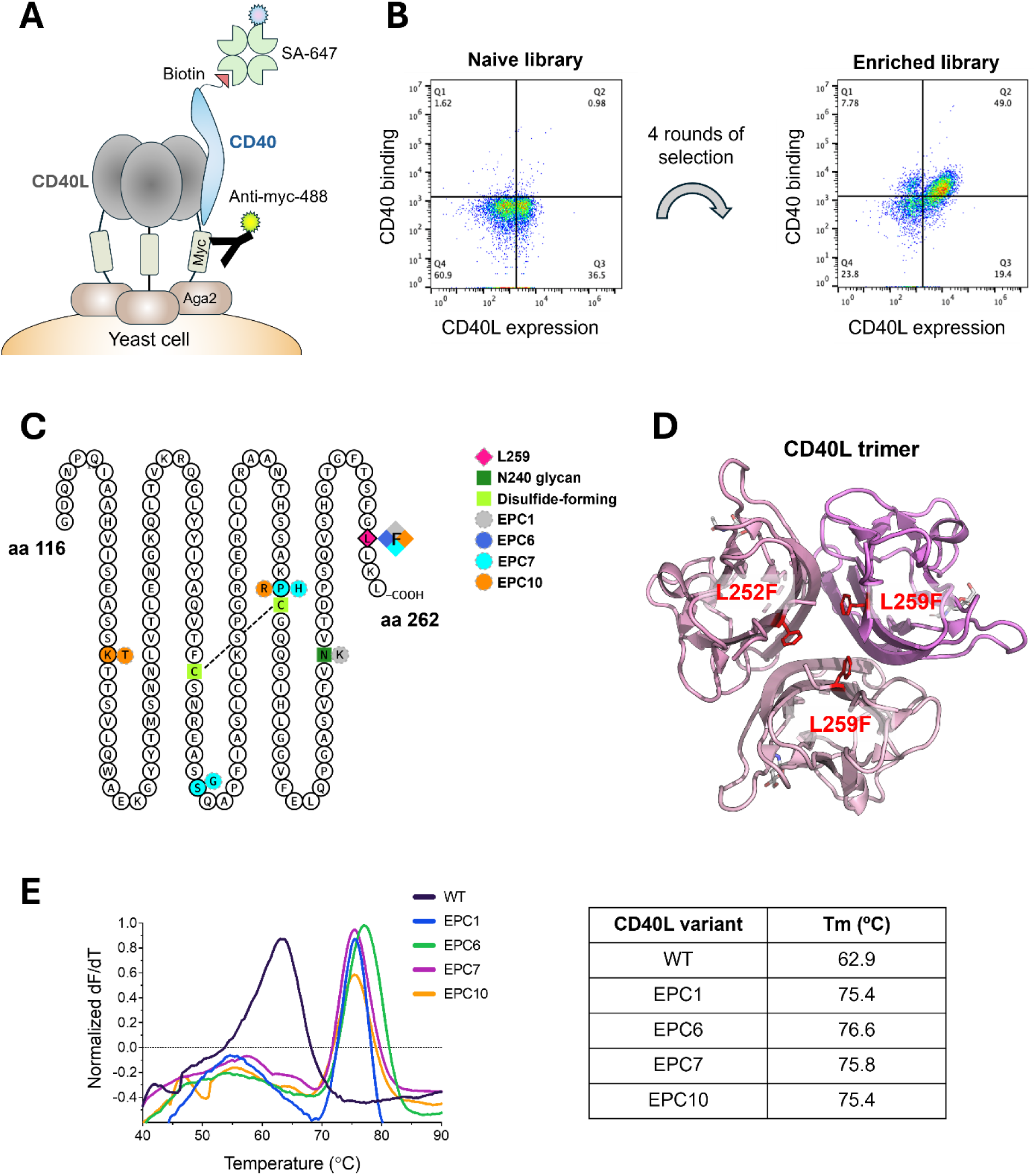
Selection and characterization of biochemically-stabilized CD40L variants. **(A)** Schematic depicting the CD40L yeast display construct. **(B)** Flow cytometry dot plots showing the enrichment of yeast capable of binding CD40 after four rounds of selection. Yeast displaying the naïve or enriched library were stained with 500 nM recombinant, biotinylated CD40 and then binding was detected with streptavidin (labeled with Alexafluor-647) and an anti-Myc antibody (labeled with Alexafluor-488). **(C)** Snake plot showing the positions and identities of amino acids that were mutated in CD40L variants. **(D)** The L259F mutation was modeled into the crystal structure of the CD40L trimer (PDB ID: 3DQ6). The mutated L259 residue is highlighted in red, and the individual protomers are colored in 3 different shades of pink. **(E)** Differential scanning fluorimetry traces (left) were used to determine the melting temperatures of purified CD40L WT and CD40L variants.

### CD40L mutants upregulate co-stimulatory markers on B cells

We next assessed the ability of EPC6 and other CD40L variants to bind and activate B cells. In a fluorescence-based assay, we found that the mutants bound to CD40 on B cells at levels comparable to the wild-type CD40L ectodomain (**Fig. 2A**, CD40L WT). Treatment of B cells with the variants also increased expression the co-stimulatory molecules CD80 and CD86, which are known to be upregulated following CD40 activation, similarly to CD40L WT (**Fig. 2B**).

**Figure 2.**
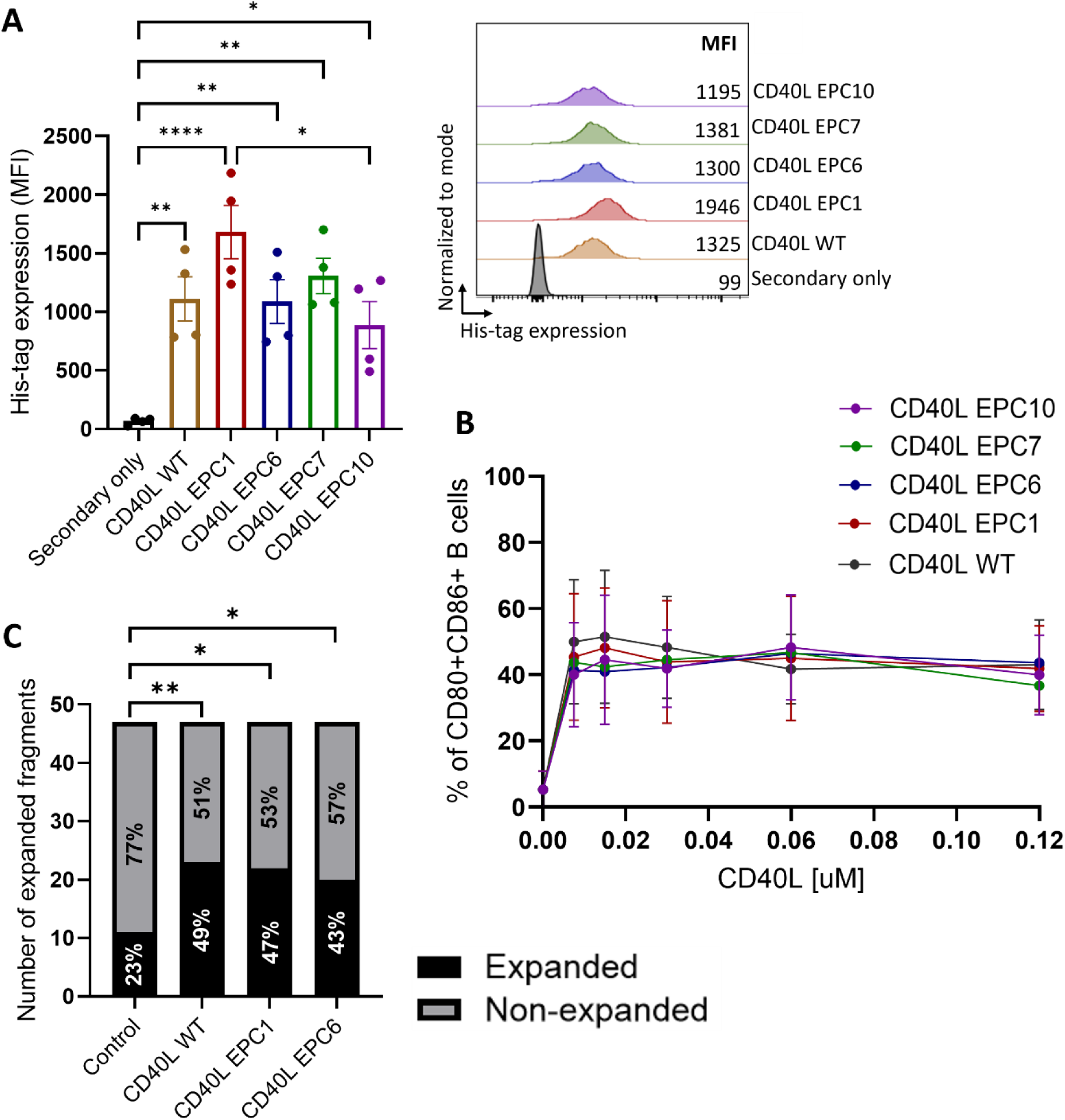
CD40L mutants promote B cell activation and improve TIL expansion. **(A)** Binding of CD40L mutants to B cells. PBMCs from healthy donors were incubated with the CD40L monomers for 20 minutes at 4°C. Secondary staining of the His tag present in the monomers was analyzed by flow cytometry on CD3⁻CD19⁺ cells, and the mean fluorescence intensity (MFI) was used as a metric of binding. Dots represent individual donors. Bars represent mean ± SEM (n = 4). A representative histogram of His-tag staining is also shown. **(B)** Dose-response analysis of B cell activation. B cells from healthy donors were cultured for 48 hours in TIL culture media or with increasing concentrations of CD40L mutants. The percentage of CD80⁺CD86⁺ cells within the B cell population was quantified by flow cytometry. Dots represent the mean result of individual donors ± SEM (n = 4). **(C)** TIL expansion analysis. Lung tumor fragments were cultured for 3–4 weeks in standard TIL expansion media (Control) or media supplemented with CD40 agonists as indicated. Shown is the TIL expansion success rate for individual tumor fragments from eight independent patients (*p ≤ 0.05; **p ≤ 0.01).

### Stimulation with CD40 agonists enhances lung TIL expansion

Addition of CD40L on the first day of TIL culture enhances TIL expansion^6^. Therefore, we tested how the L259-stabilized CD40L variants affected TIL expansion. Lung tumor fragments were cultured in standard TIL expansion media containing IL-2 alone (control) or media supplemented with CD40L WT, CD40L EPC1 or CD40L EPC6, and TIL expansion was assessed after 3–4 weeks. Across eight independent samples, TIL expansion was successful in 23% of control cultures, whereas the addition of CD40L constructs increased the success rate to 43–49% (**Fig. 2C**; *p≤0.05 and **p≤0.01). The CD40L EPC1 and CD40L EPC6 mutants enhanced TIL expansion to a similar extent to the wild-type CD40L (CD40L WT; **Fig. 2C**). Taken together, the above findings suggest that biochemically-stabilized CD40L variants retain their functional properties in B cell-and TIL-based assays.

### Engineering CD40L-41BBL fusions as synthetic immune synapses

Our previous findings support targeting tumor-infiltrating B cells as a viable strategy to improve TIL expansion and quality. In the current study, we explored whether simultaneous stimulation of tumor-infiltrating B and T cells could further optimize TIL therapy. B cell stimulation was achieved through the CD40–CD40L axis, using the CD40L^EPC6^ mutant based on its favorable biochemical properties. For T cell stimulation, we selected the 41BB–41BBL pathway, building on prior work showing that addition of anti-41BB agonistic antibodies to TIL cultures enhanced CD8⁺ T cell expansion.^9^ ^10^ ^15^ ^16^ Guided by this rationale, we designed a bi-specific molecule comprised of CD40^EPC6^ fused to a 41BBL ectodomain via a trimeric leucine zipper (**Fig. 3A**). This design was intended to create a “synthetic immune synapse” between T cells and B cells, or antigen presenting cells, and incorporates both a stabilizing mutation and trimeric scaffold to enforce native-like multimerization of the ligands.

**Figure 3.**
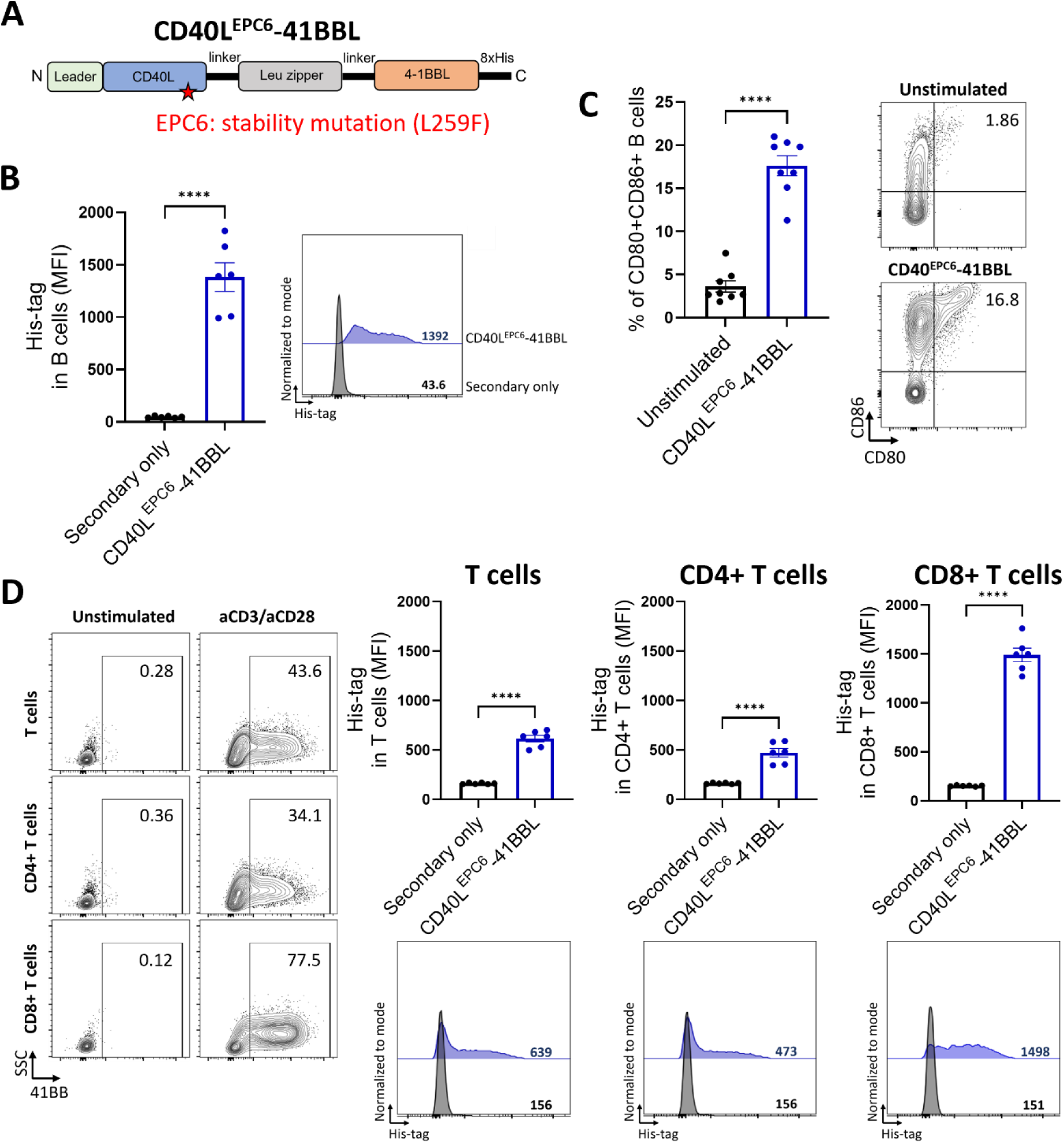
CD40L^EPC6^-41BBL binds to B and T cells and upregulates B cell activation markers. **(A)** Schematic representation of bispecific fusion protein consisting of a CD40L^EPC6^ ectodomain linked to a 41BBL ectodomain through a leucine zipper region. **(B)** Binding to B cells was quantified through secondary staining of the His-tag on CD3⁻CD19⁺ cells by flow cytometry. PBMCs from healthy donors were incubated with 0.043 µM CD40L**^EPC6^**-41BBL for 20 minutes at 4°C Dots represent individual donors. Bars represent mean ± SEM (n = 3, duplicate). A representative histogram of His-tag staining is shown. **(C)** B cell activation. B cells from healthy donors were cultured for 48 hours in control media or with 0.043 µM CD40^EPC6^-41BBL. The percentage of CD80⁺CD86⁺ cells within the B cell population (CD3⁻CD19⁺) was quantified by flow cytometry. Dots represent individual donors. Bars represent mean ± SEM (n = 4, duplicate). **(D)** Binding to T cells. T cells isolated from health donors PBMCs were stimulated with Dynabeads for 24 hours to induce 41BB expression, then incubated with CD40L^EPC6^-41BBL for 20 minutes at 4°C. CD40L^EPC6^-41BBL binding to CD4⁺ and CD8⁺ T cells were quantified through secondary staining of the His tag by flow cytometry. Dots represent individual donors. Bars represent mean ± SEM (n = 3, duplicate). Representative dot plots of 41BB expression and histograms of His-tag staining are shown.

### The CD40L^EPC6^-41BBL bispecific binds B and T cells and increases expression of co-stimulatory markers

Similar to our approach with the CD40L constructs, we evaluated whether the fusion molecule CD40L^EPC6^-41BBL could bind to and activate B cells. As shown in **Figure 3B**, CD40L^EPC6^-41BBL demonstrated significant binding to CD40 on B cells (p≤0.0001). We then examined its ability to stimulate B cells. To that end, B cells were incubated with CD40L^EPC6^-41BBL at a concentration of 0.043 µM (as previously established for CD40L alone^6^, and expression levels of CD80 and CD86 were analyzed by flow cytometry. As shown in **Figure 3C**, this intervention resulted in a 5.4-fold increase in co-expression of CD80 and CD86 on B cells (p≤0.0001). Because CD40L^EPC6^-41BBL was designed to interact with both B and T cells, we next evaluated its ability to bind activated T cells. T cells from healthy donors were stimulated with Dynabeads for 24 hours, after which binding of CD40L^EPC6^-41BBL was assessed. As shown in **Figure 3D**, CD40L^EPC6^-41BBL demonstrated significant binding to 41BBL expressed on activated T cells (p≤0.0001), with predominant binding to CD8+ T cells.

### CD40L^EPC6^-41BBL enhances CD8+ T cell expansion

As a next step, we evaluated the effect of CD40L^EPC6^-41BBL on TIL expansion. Lung tumor fragments were cultured in standard IL-2 TIL expansion media (control), or media supplemented with CD40L^EPC6^-41BBL. After 3–4 weeks, total TIL yield was greater in the CD40L^EPC6^-41BBL group compared to the control. Among nineteen independent samples, TIL expansion was successful in 38% of control cultures versus 54% with CD40L^EPC6^-41BBL (**Fig. 4A**; *p≤0.05, **p≤0.01). To complement these results, melanoma fragments were cultured under the same conditions, IL-2 only, as control, or with the addition of bi-specific protein, CD40L^EPC6^-41BBL. As shown in **Figure 4B**, a similar pattern was observed: greater TIL yield in the CD40L^EPC6^-41BBL group (p≤0.05), and improved success rates from 20% to 44% across twelve samples (p≤0.001). Consistent with previous findings using anti-41BBL agonistic antibodies^9^, CD40L^EPC6^-41BBL markedly enhanced CD8⁺ T-cell expansion in both tumor types (**Fig. 4C**; p≤0.0001).

**Figure 4.**
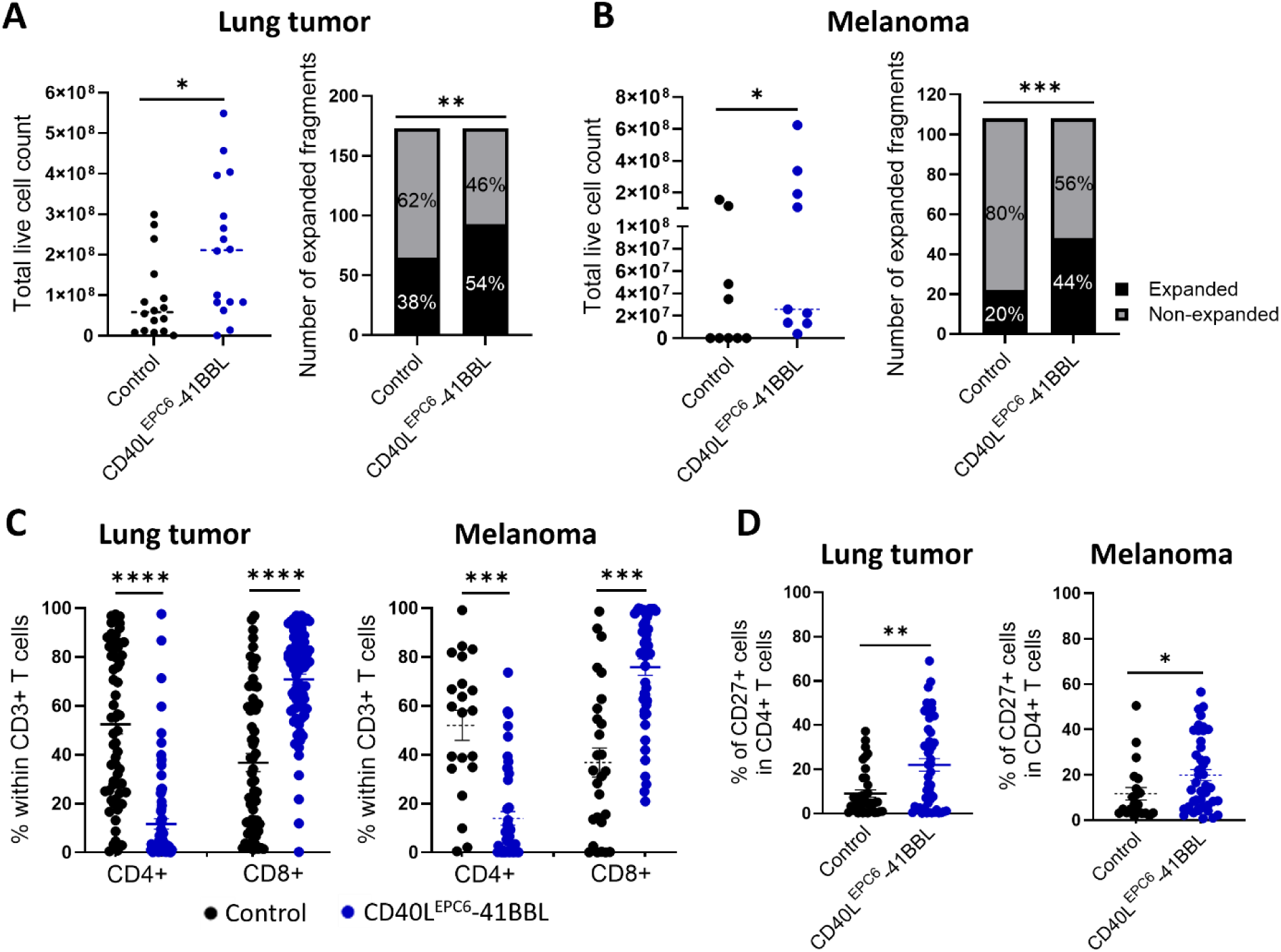
CD40^EPC6^-41BBL enhances CD8^+^ T cell expansion. **(A)** Lung tumor fragments were cultured for 3–4 weeks in standard TIL expansion media (Control) or media supplemented with 0.043 µM CD40L^EPC6^-4BBL. Total live cells and TIL expansion success rates for individual tumor fragments from nineteen independent patients are shown (*p ≤ 0.05, **p ≤ 0.01). **(B)** Melanoma fragments were cultured for 3–4 weeks in standard TIL expansion media (Control) or media supplemented with CD40L^EPC6^-4BBL. Total live cell counts and TIL expansion success rates for individual tumor fragments from twelve independent patients are shown (*p ≤ 0.05, ***p ≤ 0.001). **(C)** Percentage of CD4^+^ and CD8^+^ T cells in expanded TIL from lung tumor and melanoma fragments shown in panels A and B (****p ≤ 0.0001, ***p ≤ 0.001). **(D)** Percentage of CD27^+^ cells within the CD4^+^ T cell subset expanded from lung tumor and melanoma fragments shown in panels A and B (**p ≤ 0.01, *p ≤ 0.05).

Using high-dimensional flow cytometry analysis with FlowSOM clustering we further explored the expression of CD4, CD8, CD56, CD39, CD69, PD-1, TIM-3, CD27 and CD62L within the CD3+ T cell population (**Suppl. Fig. 1 and 2**). We identified ten clusters, six of which had the same expression pattern shared between the tumor types; **Supplemental Figures 1 and 2** describes those clusters based on a color pattern. As expected, the CD4⁺CD8^-^ T-cell clusters were enriched in the control TIL (**Suppl. Fig. 1** - Clusters 3 and 4; **Suppl. Fig. 2** - Clusters 1 and 6); CD39 expression was observed in all four clusters, meanwhile, the CD69 expression was only observed in lung tumor cluster 3 and melanoma cluster 1. No other markers expression was observed in those clusters (**Suppl. Fig. 1B, D and 2B, D)**. A variety of CD4^-^CD8^+^ T cell clusters were enriched in the CD40L^EPC6^-41BBL TIL (**Suppl. Fig. 1** - Clusters 5, 6, 7, 9 and 10; **Suppl. Fig. 2** - Clusters 4, 5, 7, 9 and 10); the expression of CD39, and the lack of expression of CD27 and PD-1 was a pattern common to all of them (**Suppl. Fig. 1B, D and 2B, D)**. The high-dimensional flow cytometry analysis also identified unconventional CD4/CD8 clusters that otherwise might have been overlooked. CD4⁻CD8⁻ clusters were identified in both tumor types (**Suppl. Fig. 1** - Clusters 1 and 2; **Suppl. Fig. 2** - Clusters 2, 3 and 8), enriched in the CD40L^EPC6^-41BBL TIL, likely representing activated T cells with downregulated lineage markers.^11^ Lung tumor cluster 1 and melanoma cluster 3 have a shared expression pattern, with expression of CD39, CD69, TIM3, and CD56, and a lack of expression of PD-1, CD62L and CD27 (**Suppl. Fig. 1B, D and 2B, D**). Notably, a unique CD4⁺CD8⁺ cluster (**Suppl. Fig. 1** - Cluster 8) appeared only in CD40L^EPC6^-41BBL-treated lung tumor samples. Interestingly it has the same expression pattern as the lung tumor cluster 1 and melanoma cluster 3 previously described.

We then went back to the conventional flow cytometry analysis to explore the CD4^+^ and CD8^+^ single subpopulations (**Suppl. Fig. 3**). No difference among the conditions was observed in the CD39/CD69, PD-1/TIM-3 and CD62L subpopulations in the CD4^+^ T cells in the lung tumor samples (**Suppl. Fig. 3A-C**). As for the melanoma CD4^+^ TIL, **Supplemental Figure 3G** demonstrates a balance between a CD39^+^CD69^+^ subpopulation increase and a CD39^-^CD69^-^ subpopulation decrease in the CD40L^EPC6^-41BBL-treated samples compared to the control. No other phenotypic difference was observed (**Suppl. Fig. 3H-I**). In the CD8^+^ T cells compartment, no clear pattern among the tumor types was observed (**Suppl. Fig. 3D-F, J-L**); compared to the control the CD40L^EPC6^-41BBL-treated lung tumor samples had a decrease in the CD39^+^CD69^-^subpopulation, meanwhile the melanoma ones had a decrease in another subpopulation, CD39^-^CD69^+^ (**Suppl. Fig. 3D and J**). For the PD-1/TIM-3 combination, the only subpopulation not altered by the different expansion conditions in the lung tumor samples, PD-1^-^TIM-3^-^ CD8+ TIL, were the ones decreased in the CD40L^EPC6^-41BBL-treated melanoma samples, combined with a small increase in the PD-1^+^TIM-3^-^ subpopulation (**Suppl. Fig. 3E and K**). The CD27 and CD62L expression in the CD8+ T cells was not altered by the CD40L^EPC6^-41BBL addition to the culture (**Suppl. Fig. 3F and L**). Interestingly, both tumor types showed an increased percentage of CD27⁺ CD4⁺ T cells in CD40L^EPC6^-41BBL cultures (**Fig. 4D**).

### CD40L^EPC6^-41BBL molecule preserves tumor reactivity of expanded TIL

To assess whether CD40L^EPC6^–41BBL influences the tumor reactivity of the resulting TIL products, expanded TIL were co-cultured with autologous tumor digests. Cultures derived from individual fragments were considered reactive based on their ability to produce IFN-γ, TNF-α, and Granzyme B, and on the inhibition of cytokine production by an anti–HLA class I blocking antibody. As shown in **Figure 5A**, lung cancer TIL showed similar rates for autologous tumor recognition in control and CD40L^EPC6^–41BBL-treated cultures, ranging between 7-11% for control TIL and 4-12% for CD40L^EPC6^–41BBL, depending on the cytokine. Interestingly, melanoma TIL showed a trend for increased tumor recognition when they were cultured in presence of CD40L^EPC6^–41BBL, ranging between 34-49% of reactive cultures, compared to 14-24% in the control group (**Fig. 5B**). These findings indicate that the superior expansion induced by CD40L^EPC6^–41BBL is not due to preferential proliferation of non-relevant bystander T cells. In fact, at least in melanoma TIL, this intervention may result in an increase of tumor-reactive clones.

**Figure 5.**
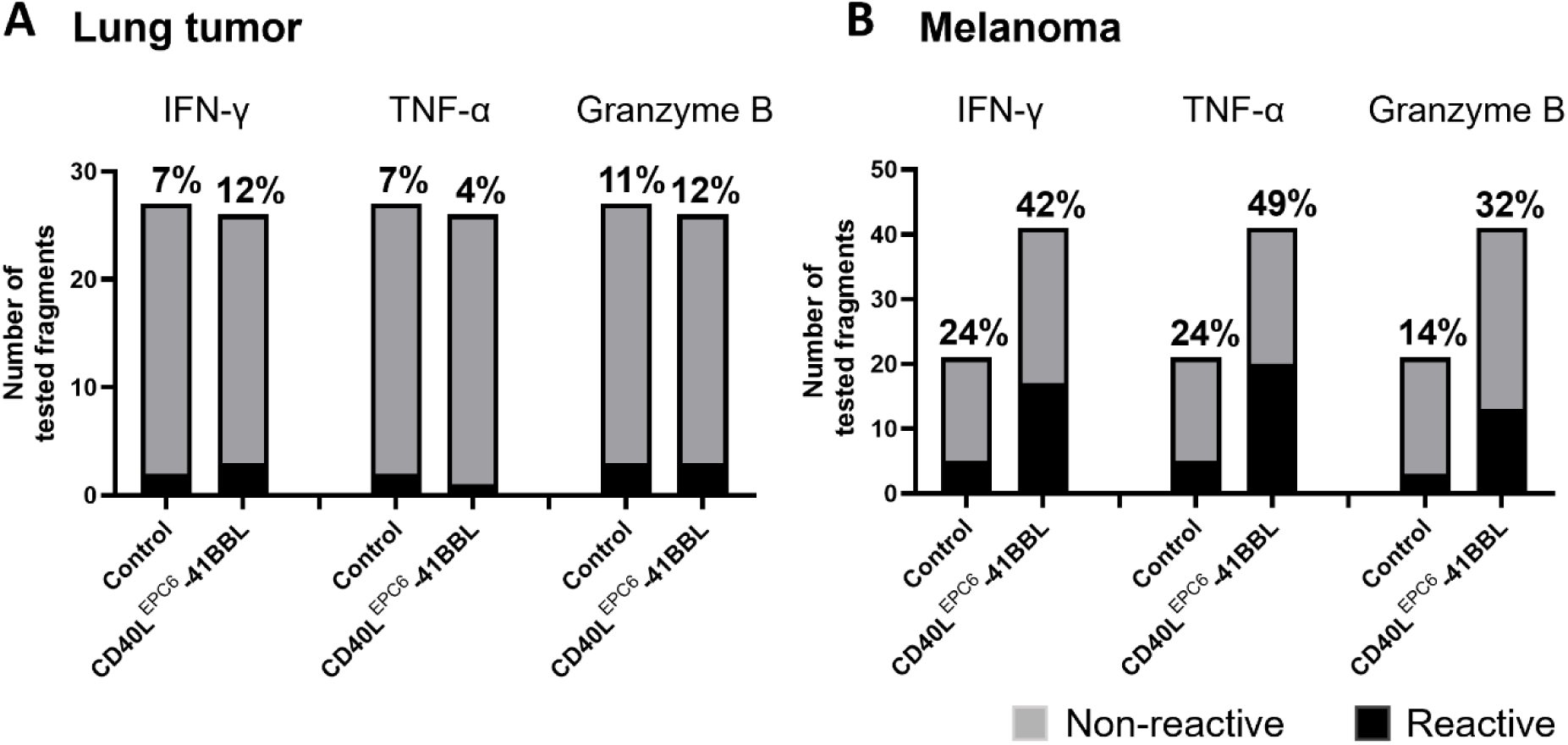
CD40^EPC6^-41BBL preserves tumor reactivity of expanded TIL. Expanded TIL from lung tumors **(A)** and melanomas **(B)** were co-cultured with autologous tumor cells for 24 hours at a 1:1 ratio. Tumor reactivity was assessed by measuring IFN-γ, TNF-α, and Granzyme B levels in the supernatant. For lung tumor samples **(A)**, 27 control fragments and 26 CD40L^EPC6^–41BBL–treated fragments were analyzed; for melanoma samples **(B)**, 21 control fragments and 41 CD40L^EPC6^–41BBL–treated fragments were evaluated. Black bars indicate the percentage of fragments classified as tumor-reactive, while gray bars represent non-reactive fragments.

## DISCUSSION

In this study, we sought to improve TIL manufacturing by simultaneously targeting two key cellular components of the tumor microenvironment: tumor-infiltrating B cells and T cells. Our findings demonstrate that dual stimulation of the CD40 and 41BB pathways using a novel bi-specific molecule, CD40L^EPC6^–41BBL, enhances TIL expansion and promotes the enrichment of CD8⁺ T cells in both lung tumor and melanoma cultures. The rationale for combining CD40 and 41BB agonism was based on complementary mechanisms of action. CD40 signaling activates B cells, increasing their antigen-presenting capacity and expression of co-stimulatory molecules^17^, whereas 41BB signaling directly promotes T cell proliferation, long-term survival, and effector function.^18^ Previous studies from our group independently demonstrated the benefits of targeting each pathway during TIL culture.^6^ ^9^ The present work extends these observations by showing that concurrent activation of both pathways further improves TIL manufacturing. Compared with standard IL-2 culture conditions, supplementation with CD40L^EPC6^–41BBL increased both total TIL yield and the frequency of successful expansions in lung tumor and melanoma patients’ samples.

Our CD40L^EPC6^ variant incorporates a stabilizing L259F mutation at the trimer interface. Although L259F was identified solely through selection for CD40 binding and increased surface expression on yeast, it also produced a marked increase in the thermal stability of the purified soluble ectodomain. One possible explanation is that this substitution stabilizes the native trimer interface, thereby increasing the fraction of properly assembled ligand during biosynthesis and secretion. Membrane-bound CD40L normally exists as a type II transmembrane protein in which the transmembrane helix and extracellular stalk constrain the spatial organization of the TNF homology domains prior to trimer formation. By strengthening hydrophobic packing within the trimer interface, L259F may partially compensate for the absence of these native stabilizing elements, promoting productive folding without measurably altering receptor signaling. The emergence of a thermostabilizing mutation through functional selection is also consistent with previous protein engineering studies in which directed evolution enriched variants with enhanced conformational stability. In many systems, including antibodies^19^, surface ligands^20^, and viral glycoproteins^21^, mutations that improve folding efficiency or stabilize the native state have been recovered despite selection being performed exclusively for ligand binding or functional activity. Our findings suggest that yeast display-based evolution of TNF superfamily ligands may similarly enrich mutations that improve both biophysical stability and manufacturability while preserving biological function.

An important observation was the preferential expansion of CD8⁺ T cells induced by CD40L^EPC6^–41BBL, since CD8⁺ TIL have been associated with favorable responses to TIL therapy.^22^ The increase in CD8⁺ T cells observed here is consistent with our previous report demonstrating the capacity of 41BB signaling to selectively support cytotoxic T cells.^9^ ^10^ Importantly, the enrichment of CD8⁺ cells was observed in both evaluated tumor types, suggesting that the effects of CD40L^EPC6^–41BBL may be broadly applicable across distinct tumor microenvironments. Even though the proportion of CD4^+^ T cells was reduced compared to the CD8^+^ T cells, the CD40L^EPC6^–41BBL treated cultures contained higher frequencies of CD27⁺ CD4⁺ T cells, a phenotype frequently associated with less differentiated T cells^23^ possessing greater proliferative potential and persistence.^24^ ^25^

The high-dimensional flow cytometry analysis provided additional insights into the phenotypic changes induced by dual stimulation. FlowSOM analysis identified several CD8⁺ T cell clusters that were consistently enriched in cultures treated with CD40L^EPC6^–41BBL. These populations expressed markers associated with activation and tumor antigen experience, including CD39 and CD69.^26^ The enrichment of these populations therefore suggests that CD40L^EPC6^–41BBL may preferentially support expansion of lymphocytes with prior tumor recognition. Our reactivity data supports this statement. Although the limited availability of autologous tumor material for co-culture, combined with the small number of patient samples, constrained our ability to detect statistically significant differences, we observed a clear trend for increased tumor reactivity in melanoma TIL expanded with CD40L^EPC6^-41BBL.

The high-dimensional flow cytometry analysis also identified CD4⁻CD8⁻ T-cell populations enriched in cultures treated with CD40L^EPC6^-41BBL. These cells expressed activation-associated markers, including CD39, and CD69, suggesting that they may represent highly activated T cells that have downregulated lineage markers during prolonged culture. However, the cellular origin and functional role of this relatively rare T cell subset is not well established. Previous studies have reported both tumor-promoting and tumor-suppressive functions for CD4⁻CD8⁻ T cells, depending on the tumor type and microenvironmental context.^27^ ^28^ Notably, Wu et al. reported that CD69 expression by CD4⁻CD8⁻ T cells was associated with anti-tumor activity, raising the possibility that the CD69⁺ populations identified in our cultures may contribute to anti-tumor immune responses. Another unique population was detected, CD4⁺CD8⁺ T cells, this time exclusively in lung tumor cultures supplemented with CD40L^EPC6^–41BBL. Although their role in adoptive cell therapy remains poorly understood, CD4⁺CD8⁺ peripheral T cells have been reported in several human cancers^29^ and intratumoral double positive cells have been described as enriched for clonally-expanded cytotoxic T cells with the ability of recognizing target cells through both class I and class II HLA molecules.^30^ Studies have suggested that this population may represent a distinct subset of tumor-reactive lymphocytes characterized by antigen-driven expansion based on clonal T cell receptor repertoires, memory-associated phenotypes, and cytotoxic activity against autologous tumor cells.^29^ ^31^

In conclusion, this study demonstrates that simultaneous activation of CD40 and 41BB signaling through a novel CD40L^EPC6^–41BBL bi-specific molecule enhances ex vivo TIL expansion and promotes enrichment of CD8⁺ T cell populations. This approach has the potential of maximizing TIL expansion from tumors with low basal lymphocytic infiltration. Importantly, it may reduce the amount of starting material needed for TIL manufacture, allowing for new protocols requiring minimally invasive procedures to procure the starting tumor material, as discussed in a co-submitted manuscript (Karapetyan et al, under review). Two ongoing clinical trials are underway to test the feasibility of this approach, NCT05681780 for patients with lung cancer and NCT06961357 for patients with melanoma.

## ACKNOWLEDGMENTS

This work was partially supported by the Suzi Q Foundation, the Moffitt Melanoma Center of Excellence, the Shultz Family Foundation, the Dr. Miriam and Sheldon G. Adelson Medical Research Foundation, and the Mark Foundation ASPIRE program. This work has been supported in part by the Cell Therapies, Flow Cytometry, Molecular Genomics and Tissue Core Facilities at the Moffitt Cancer Center, an NCI designated Comprehensive Cancer Center (P30-CA076292). The authors would like to thank the Tissue Core Facility team for their assistance in coordinating the collection of fresh specimens.

## AUTHORS CONTRIBUTIONS

Conception and design: DAD, PH, VL, RAMR. Study supervision: DAD. Development and methodology execution: RAMR, MSB, JC, JRA, EMP. Yeast display selections: KH. Construct design and protein purification: AAR and MGG. Analysis and interpretation of data: RAMR, MSB, SPT, DAD. Scientific advisors: LK, BC, SPT, PH, VL. Sample collection support: JA, LK, BC. Writing the manuscript: RAMR, VL, DAD. All authors reviewed and approved the final manuscript.

## REFERENCES

1. Maiocco G, Ernani V, Gustafson MP, et al. Cellular Therapies in Solid Tumors: Are We Ready for Primetime? JCO Oncol Pract 2026:OP2600119. doi: 10.1200/OP-26-00119 [published Online First: 20260520]

2. Rosenberg SA, Restifo NP, Yang JC, et al. Adoptive cell transfer: a clinical path to effective cancer immunotherapy. Nat Rev Cancer 2008;8(4):299–308. doi: 10.1038/nrc2355

3. Rizkallah J, Wehbeh BED, Wassouf W, et al. Adoptive cell therapies in solid tumors: current clinical landscape, challenges, and future directions. Front Immunol 2026;17:1800292. doi: 10.3389/fimmu.2026.1800292 [published Online First: 20260514]

4. Chen R, Johnson J, Rezazadeh A, et al. Tumour-infiltrating lymphocyte therapy landscape: prospects and challenges. BMJ Oncol 2025;4(1):e000566. doi: 10.1136/bmjonc-2024-000566 [published Online First: 20250804]

5. Pherez-Farah A, Boncompagni G, Chudnovskiy A, et al. The Bidirectional Interplay between T Cell-Based Immunotherapies and the Tumor Microenvironment. Cancer Immunol Res 2025;13(4):463–75. doi: 10.1158/2326-6066.CIR-24-0857

6. Rossetti RAM, Tordesillas L, Beatty MS, et al. CD40L stimulates tumor-infiltrating B-cells and improves ex vivo TIL expansion. J Immunother Cancer 2025;13(4) doi: 10.1136/jitc-2024-011066 [published Online First: 20250408]

7. Smith AS, Knochelmann HM, Wyatt MM, et al. B cells imprint adoptively transferred CD8(+) T cells with enhanced tumor immunity. J Immunother Cancer 2022;10(1) doi: 10.1136/jitc-2021-003078

8. Ruffin AT, Toliopoulos V, Smith AS, et al. TLR9 Agonists Potentiate Adoptive T Cell Therapy in Cancer through a B Cell-CD2 Costimulatory Axis. Cancer Res 2026 doi: 10.1158/0008-5472.CAN-26-0202 [published Online First: 20260706]

9. Innamarato P, Asby S, Morse J, et al. Intratumoral Activation of 41BB Costimulatory Signals Enhances CD8 T Cell Expansion and Modulates Tumor-Infiltrating Myeloid Cells. J Immunol 2020;205(10):2893–904. doi: 10.4049/jimmunol.2000759 [published Online First: 20201005]

10. Chacon JA, Sarnaik AA, Chen JQ, et al. Manipulating the tumor microenvironment ex vivo for enhanced expansion of tumor-infiltrating lymphocytes for adoptive cell therapy. Clin Cancer Res 2015;21(3):611–21. doi: 10.1158/1078-0432.CCR-14-1934 [published Online First: 20141203]

11. Feldhaus MJ, Siegel RW, Opresko LK, et al. Flow-cytometric isolation of human antibodies from a nonimmune Saccharomyces cerevisiae surface display library. Nat Biotechnol 2003;21(2):163–70. doi: 10.1038/nbt785 [published Online First: 20030121]

12. Ming Q, Antfolk D, Price DA, et al. Structural basis for mouse LAG3 interactions with the MHC class II molecule I-A(b). Nat Commun 2024;15(1):7513. doi: 10.1038/s41467-024-51930-5 [published Online First: 20240829]

13. Naito M, Hainz U, Burkhardt UE, et al. CD40L-Tri, a novel formulation of recombinant human CD40L that effectively activates B cells. Cancer Immunol Immunother 2013;62(2):347–57. doi: 10.1007/s00262-012-1331-4 [published Online First: 20120825]

14. An HJ, Kim YJ, Song DH, et al. Crystallographic and mutational analysis of the CD40-CD154 complex and its implications for receptor activation. J Biol Chem 2011;286(13):11226–35. doi: 10.1074/jbc.M110.208215 [published Online First: 20110201]

15. Chacon JA, Sarnaik AA, Pilon-Thomas S, et al. Triggering co-stimulation directly in melanoma tumor fragments drives CD8(+) tumor-infiltrating lymphocyte expansion with improved effector-memory properties. Oncoimmunology 2015;4(12):e1040219. doi: 10.1080/2162402X.2015.1040219 [published Online First: 20150701]

16. Chacon JA, Wu RC, Sukhumalchandra P, et al. Co-stimulation through 4-1BB/CD137 improves the expansion and function of CD8(+) melanoma tumor-infiltrating lymphocytes for adoptive T-cell therapy. PLoS One 2013;8(4):e60031. doi: 10.1371/journal.pone.0060031 [published Online First: 20130401]

17. Possamai D, Page G, Panes R, et al. CD40L-Stimulated B Lymphocytes Are Polarized toward APC Functions after Exposure to IL-4 and IL-21. J Immunol 2021;207(1):77–89. doi: 10.4049/jimmunol.2001173 [published Online First: 20210616]

18. Bagheri S, Safaie Qamsari E, Yousefi M, et al. Targeting the 4-1BB costimulatory molecule through single chain antibodies promotes the human T-cell response. Cell Mol Biol Lett 2020;25:28. doi: 10.1186/s11658-020-00219-8 [published Online First: 20200422]

19. Famm K, Hansen L, Christ D, et al. Thermodynamically stable aggregation-resistant antibody domains through directed evolution. J Mol Biol 2008;376(4):926–31. doi: 10.1016/j.jmb.2007.10.075 [published Online First: 20071104]

20. Gonzalez-Perez D, Das S, Antfolk D, et al. Affinity-matured DLL4 ligands as broad-spectrum modulators of Notch signaling. Nat Chem Biol 2023;19(1):9–17. doi: 10.1038/s41589-022-01113-4 [published Online First: 20220901]

21. McLellan JS, Chen M, Joyce MG, et al. Structure-based design of a fusion glycoprotein vaccine for respiratory syncytial virus. Science 2013;342(6158):592–8. doi: 10.1126/science.1243283

22. Granhoj JS, Mannering SMH, Madsen K, et al. Phenotypic and Functional Characteristics of CD8+ T cells Predict Clinical Outcome Following TIL Therapy in a Randomized Phase III Trial in Advanced Melanoma. Clin Cancer Res 2026 doi: 10.1158/1078-0432.CCR-25-4097 [published Online First: 20260518]

23. Tran KQ, Zhou J, Durflinger KH, et al. Minimally cultured tumor-infiltrating lymphocytes display optimal characteristics for adoptive cell therapy. J Immunother 2008;31(8):742–51. doi: 10.1097/CJI.0b013e31818403d5

24. Hendriks J, Xiao Y, Borst J. CD27 promotes survival of activated T cells and complements CD28 in generation and establishment of the effector T cell pool. J Exp Med 2003;198(9):1369–80. doi: 10.1084/jem.20030916 [published Online First: 20031027]

25. Remedios KA, Meyer L, Zirak B, et al. CD27 Promotes CD4(+) Effector T Cell Survival in Response to Tissue Self-Antigen. J Immunol 2019;203(3):639–46. doi: 10.4049/jimmunol.1900288 [published Online First: 20190617]

26. Mullinax JE, Hall M, Beatty M, et al. Expanded Tumor-infiltrating Lymphocytes From Soft Tissue Sarcoma Have Tumor-specific Function. J Immunother 2021;44(2):63–70. doi: 10.1097/CJI.0000000000000355

27. Wu Z, Zheng Y, Sheng J, et al. CD3(+)CD4(-)CD8(-) (Double-Negative) T Cells in Inflammation, Immune Disorders and Cancer. Front Immunol 2022;13:816005. doi: 10.3389/fimmu.2022.816005 [published Online First: 20220210]

28. Liu XF, Song B, Sun CB, et al. Tumor-infiltrated double-negative regulatory T cells predict outcome of T cell-based immunotherapy in nasopharyngeal carcinoma. Cell Rep Med 2025;6(5):102096. doi: 10.1016/j.xcrm.2025.102096 [published Online First: 20250501]

29. Schad SE, Chow A, Mangarin L, et al. Tumor-induced double positive T cells display distinct lineage commitment mechanisms and functions. J Exp Med 2022;219(6) doi: 10.1084/jem.20212169 [published Online First: 20220523]

30. Li T, Ilano A, Arias-Badia M, et al. Functionally heterogeneous intratumoral CD4(+)CD8(+) double-positive T cells can give rise to single-positive T cells. Proc Natl Acad Sci U S A 2026;123(4):e2506168123. doi: 10.1073/pnas.2506168123 [published Online First: 20260120]

31. Menard LC, Fischer P, Kakrecha B, et al. Renal Cell Carcinoma (RCC) Tumors Display Large Expansion of Double Positive (DP) CD4+CD8+ T Cells With Expression of Exhaustion Markers. Front Immunol 2018;9:2728. doi: 10.3389/fimmu.2018.02728 [published Online First: 20181126]

