## Supplemental Material for "Dual stimulation of CD40 and 41BB pathways during ex-vivo TIL expansion enhances CD8+ T cell expansion"

**Supplemental Table 1.** Antibodies for the flow cytometry assays.

| Antibody | Fluorochrome | Clone | Titer | Manufacturer | Catalog # |
|---|---|---|---|---|---|
| **41BB** | BV421 | 4B4-1 | 1:50 | Biolegend | 309820 |
| **CD3** | BUV395 | SK7 | 1:100 | BD | 564001 |
| **His-Tag** | Alexa Fluor 647 | - | 1:50 | Cell Signaling | 14931S |
| **CD4** | BUV496 | SK3 | 1:100 | BD | 612936 |
| **CD4** | BV785 | OKT4 | 1:25 | Biolegend | 317442 |
| **CD8** | PECy7 | RPA-T8 | 1:400 | BD | 557746 |
| **CD19** | BV605 | HIB19 | 1:100 | Biolegend | 302244 |
| **CD27** | BV605 | O323 | 1:100 | Biolegend | 302830 |
| **CD39** | FITC | A1 | 1:200 | Biolegend | 328206 |
| **CD40** | BV786 | 5C3 | 1:100 | Biolegend | 334340 |
| **CD56** | PE | NCAM-1 | 1:50 | BD | 555516 |
| **CD62L** | BV421 | DREG-56 | 1:50 | Biolegend | 304828 |
| **CD69** | APC | FN50 | 1:25 | BD | 555533 |
| **CD80** | BV650 | 2D10 | 1:25 | Biolegend | 305227 |
| **CD86** | PECy7 | IT2.2 | 1:400 | Biolegend | 305422 |
| **HLA-DR** | FITC | Tu39 | 1:400 | BD | 555558 |
| **PD-1** | Alexa Fluor 700 | EH12.2H7 | 1:50 | Biolegend | 329952 |
| **TIM-3** | BV650 | 7D3 | 1:100 | BD | 565564 |
| **Live/Dead** | Near IR | - | 1:500 | Invitrogen | L10119 |

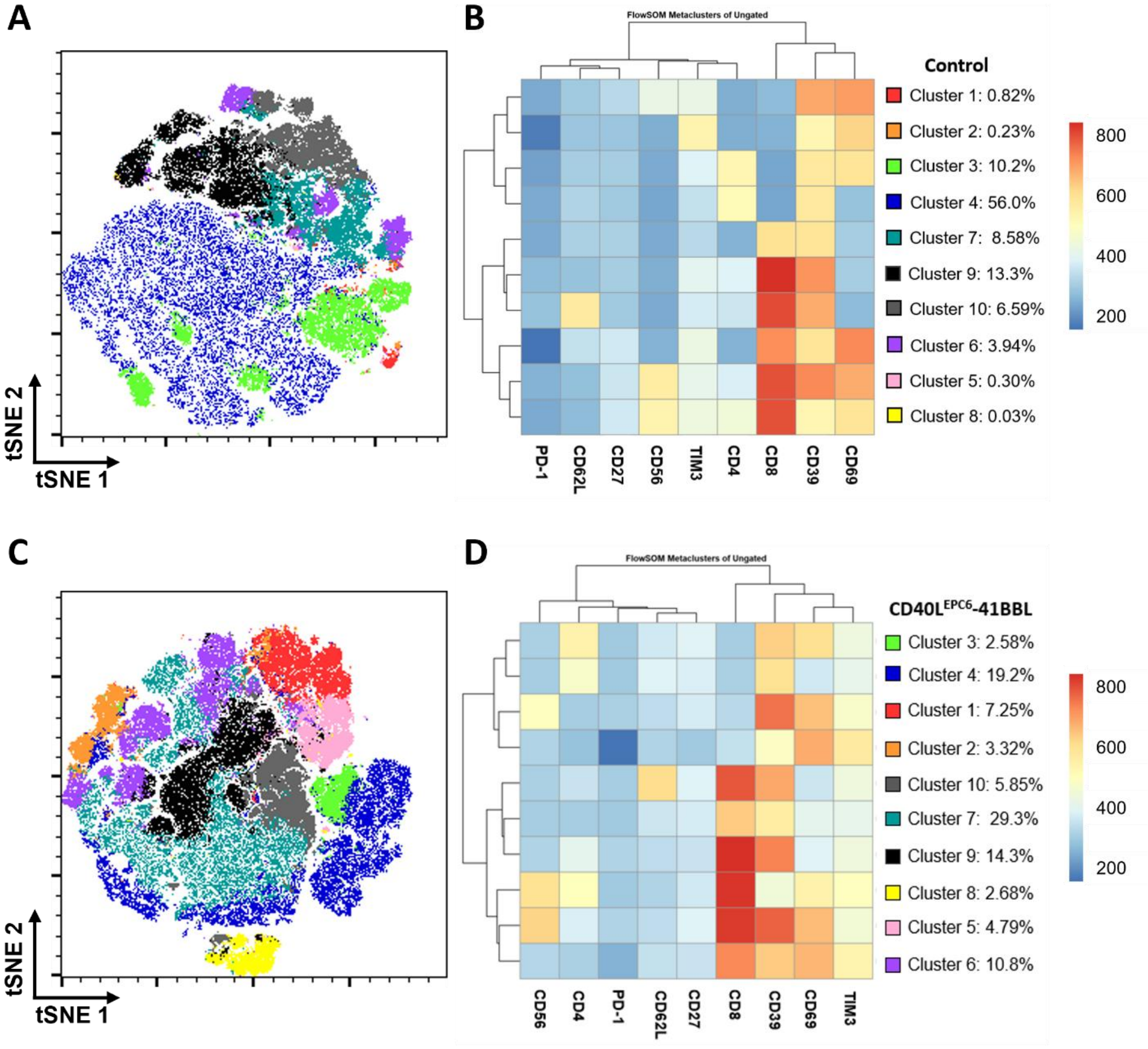


**E**

| | |
|---|---|
| **CD4- CD8-** | Clusters: **1**, 2. |
| **CD4+ CD8-** | Clusters: **3**, **4**. |
| **CD4- CD8+** | Clusters: **5**, **6**, 7, **9**, 10. |
| **CD4+ CD8+** | Clusters: 8. |

**Supplemental Figure 1. High-dimensional flow cytometry analysis with FlowSOM clustering in lung tumor samples.** FlowSOM-generated tSNE maps and metacluster heatmaps for each culture condition. Lung tumor fragments were cultured for 3–4 weeks in **(A-B)** standard TIL expansion media (Control, IL-2 only) or **(C-D)** media supplemented with 0.043 uM CD40L$^{EPC6}$-4BBL (n=10). **(E)** CD4 and CD8 expression cluster distribution.

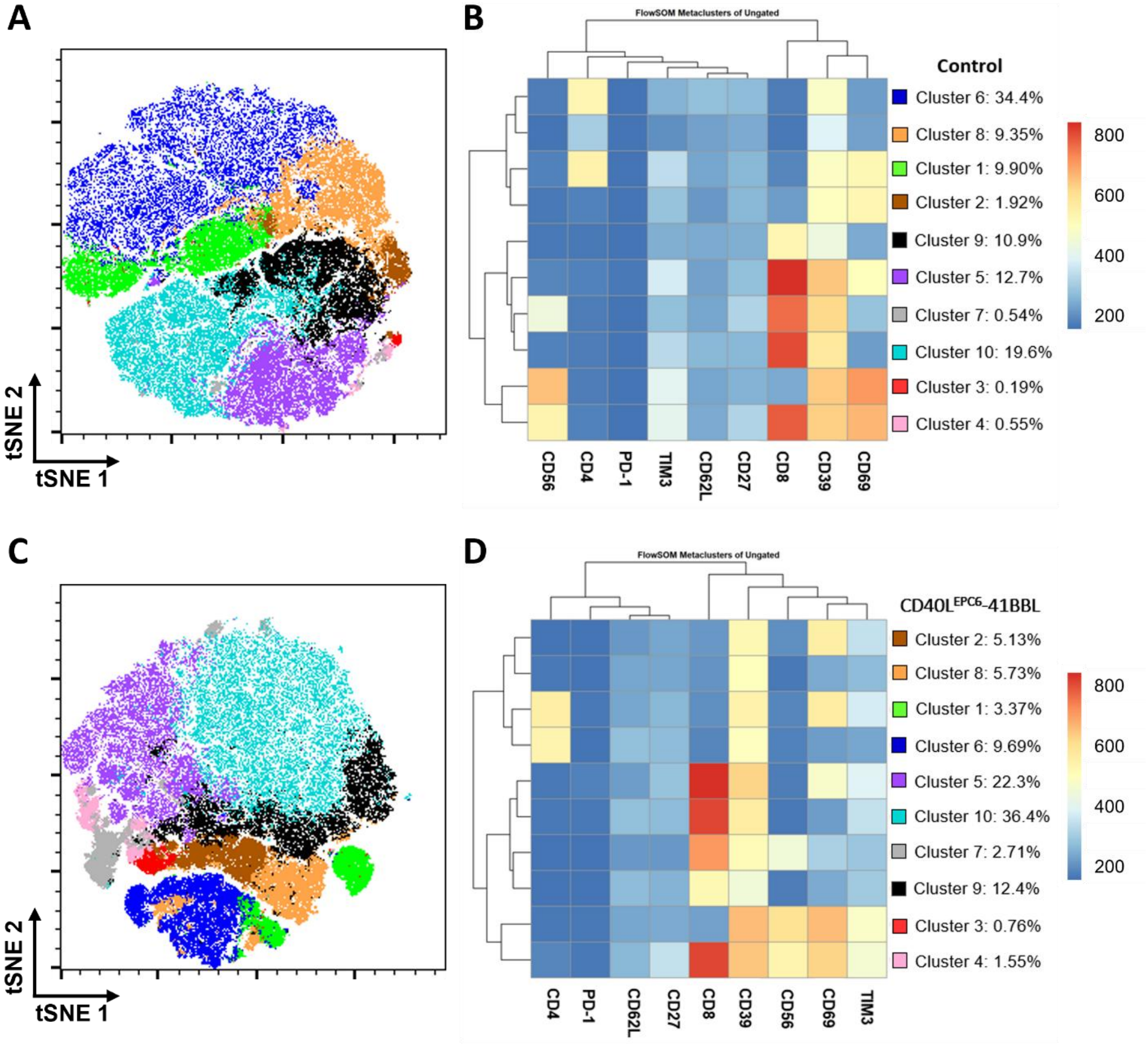


**E**

| | |
|---|---|
| **CD4- CD8-** | Clusters: 2, **3**, 8. |
| **CD4+ CD8-** | Clusters: **1**, **6.** |
| **CD4- CD8+** | Clusters: **4**, **5**, 7, **9**, 10. |
| **CD4+ CD8+** | Clusters: none. |

**Supplemental Figure 2. High-dimensional flow cytometry analysis with FlowSOM clustering in melanoma samples.** FlowSOM-generated tSNE maps and metacluster heatmaps for each culture condition. TIL. Melanoma fragments were cultured for 3–4 weeks in **(A-B)** standard TIL expansion media (Control, IL-2 only) or **(C-D)** media supplemented with 0.043 uM CD40L$^{EPC6}$-4BBL (n=9). **(E)** CD4 and CD8 expression cluster distribution.

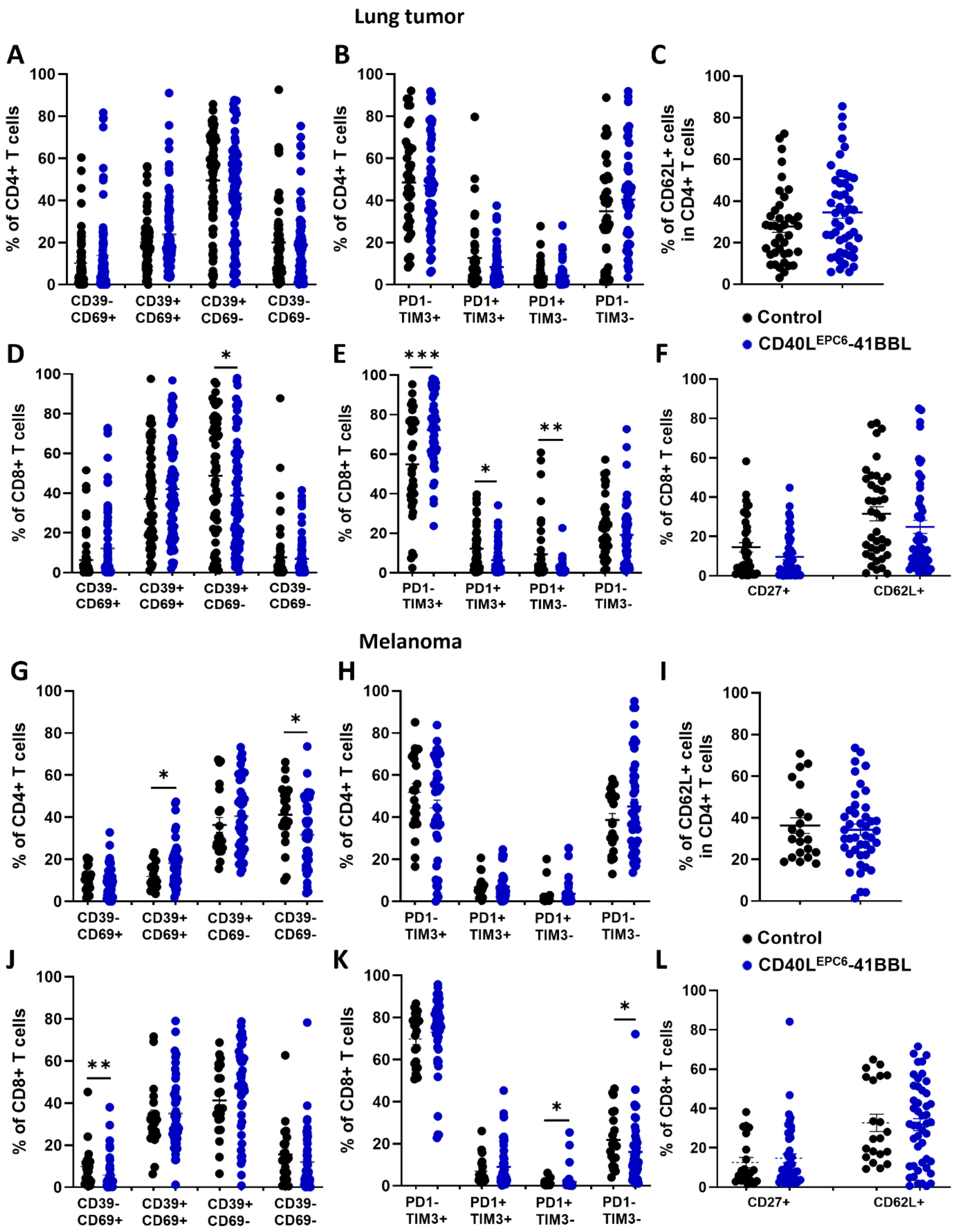


**Supplemental Figure 3. Conventional flow cytometry analysis in lung tumor and melanoma samples.** Fragments were cultured for 3–4 weeks in standard TIL expansion media (Control) or media supplemented with 0.043 μM CD40L$^{EPC6}$-4BBL **(A-F)** Percentage of CD4$^{+}$ and CD8$^{+}$ T cells subpopulations in expanded TIL from lung tumor fragments. **(G-L)** Percentage of CD4$^{+}$ and CD8$^{+}$ T cells subpopulations in expanded TIL from melanoma fragments. *p ≤ 0.05, **p ≤ 0.01, ***p ≤ 0.001.
